# Waking Up in the Dream Lab: A Lab-Based Lucid Dream Induction Paradigm Using Virtual Reality and Sensory Stimulation

**DOI:** 10.64898/2026.03.11.711049

**Authors:** Emma Peters, Johanna Heitmann, Nicolas Morath, Melanie Roth, Nicole Bühler, Eva Nussbaumer, Xinlin Wang, Ralf Kredel, Simon Maurer, Martin Dresler, Daniel Erlacher

**Author notes:** Corresponding author: Emma Peters.

## Abstract

Lucid dreaming (LD), during which the dreamer is aware that they are dreaming, is frequently induced in laboratory settings by delivering sensory cues during rapid eye movement (REM) sleep. These cues should be incorporated into ongoing dreams and can trigger reflective awareness. This approach relies on the continuity between waking experiences and dream content. In sleep laboratories, participants often dream of the experimental setting itself (lab dreaming), providing a predictable context in which lucidity may emerge. The present studies leveraged this phenomenon by explicitly training participants to associate the sleep laboratory with reflective awareness prior to sleep. Across three studies (total N = 101), participants completed a morning nap following verbal LD instructions and presleep audio designed to prime recognition of the laboratory context in dreams. In addition, conditions included immersive virtual reality (VR) rehearsal of the laboratory environment, VR combined with haptic stimulation (HS) during REM sleep, or VR containing subtle fake system errors intended to prompt reflective checking. LD frequency was assessed through external ratings of signal-verified LD (SVLD) dream reports. Lucidity rates were high across all conditions, with approximately 40-45% of dreams externally rated as lucid and 11%-32% SVLDs occurring in every group. However, neither VR rehearsal, haptic stimulation, nor implicit VR errors increased lucidity relative to the baseline laboratory induction procedure. Exploratory analyses investigated the overlap between laboratory dreaming, false awakenings (FAs), and lucidity. These findings suggest that explicit training focused on the predictable context of the sleep laboratory may already provide a powerful pathway to lucidity, with additional technological manipulations offering limited benefit under a single-nap protocol.

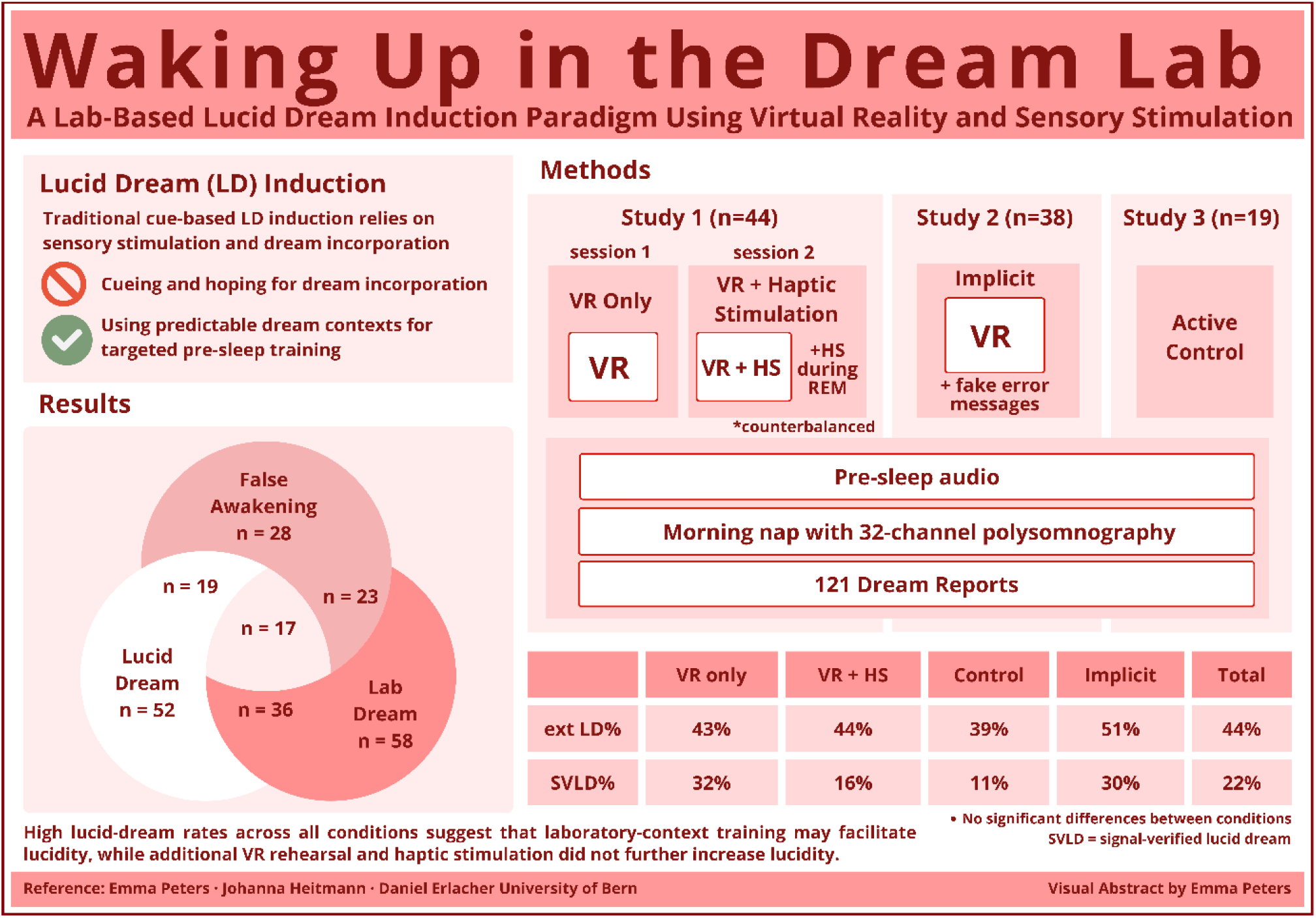

## 1. Introduction

Lucid dreaming (LD) is a unique state in which a person becomes aware that they are dreaming while the dream continues (La Berge, 1985). It has long fascinated sleep and consciousness researchers, as it allows direct access to metacognitive processing within sleep (Voss et al., 2009; Windt & Metzinger, 2007), and has been applied across diverse domains, including nightmare therapy (Ouchene et al., 2023), motor skill rehearsal, creativity enhancement, problem-solving (Stumbrys & Daniels, 2010), self-exploration (Schädlich & Erlacher, 2012), and motor practice (Bonamino et al., 2023; Peters et al., 2023). Numerous induction techniques have been proposed to elicit lucidity, including cognitive strategies such as reality testing, mnemonic training, and wake-back-to-bed protocols (Tan & Fan, 2022), as well as external stimulation methods that aim to deliver cues during rapid eye movement (REM) sleep, such as light flashes, sounds, or tactile vibrations (Carr et al., 2020; Salvesen et al., 2024).

A promising complementary approach builds on the principle of dream continuity (Schredl, 2006; Wamsley & Stickgold, 2009), and specifically, dreaming in the sleep laboratory. In lab dreams, participants dream of the experimental environment, polysomnography (PSG) setup, strange or broken equipment, or the experimenters themselves (Picard-Deland et al., 2021) and can experience false awakenings (FA). FAs are highly realistic dreams of waking up (Green, 1990) and are among the most common forms of lab-incorporated dreaming (Buzzi, 2019). Because lab dreams occur reliably in sleep studies, they provide a predictable context for LD induction. Instead of trying to cue specific dream content, we can use the anticipated sleep-lab setting to tailor waking LD training. If participants are trained to critically reflect on their state when encountering the laboratory environment, recognizing this familiar context in a dream may facilitate the transition to lucid awareness.

Virtual reality (VR) provides a controlled, immersive way to rehearse specific environments before sleep, potentially strengthening wake–dream continuity. VR experiences can be incorporated into subsequent dreams (Morris et al., 2025; Picard-Deland et al., 2020) and allow for the controlled introduction of subtle, dream-like prediction violations while maintaining spatial presence (Denzer et al., 2022). Gott et al. (2020) systematically tested VR-based LD induction using a university cafeteria-themed VR environment (VRE) with mild bizarre elements (e.g., malfunctioning clocks, altered gravity, characters staring). Participants performed metacognitive reality checks during twelve 15-minute training sessions, increasing dream lucidity relative to a passive control group but not relative to an active control group practicing standard induction techniques (reality checks and dream journaling).

Although numerous induction techniques have been proposed, few approaches explicitly leverage predictable dream contexts. Laboratory dreams and FAs occur with notable frequency in sleep studies, yet existing induction methods rarely use this stable context as a training target. Similarly, VR has primarily been used to rehearse generic reality-checking strategies rather than to prime reflective awareness within a realistic simulation of the sleep laboratory itself. The present study, therefore, investigates whether explicitly training participants to recognize the laboratory context as a potential dream setting can facilitate lucid-dream induction.

To test this idea, we developed a VRE of the University of Bern sleep laboratory. Participants completed a brief VR rehearsal prior to sleep in which they explored the simulated lab environment and practiced reality checks. The VR task was designed to strengthen the association between encountering the laboratory setting and engaging in reflective awareness. In one condition, VR training was combined with vibro-tactile stimulation during REM sleep, and in another with subtle fake system-error messages embedded in the VR environment. Overtly surreal VR events are easily interpreted as part of the simulation and therefore rarely evoke genuine reflection. In contrast, subtle, realistic system glitches and bugs create momentary ambiguity about whether the disruption originates in the VR scenario or the actual laboratory. Such minor inconsistencies were intended to mimic the kinds of ambiguities often reported in laboratory dreams and FAs, which may prompt metacognitive reflection.

The primary aim of this study was to test whether immersive VR rehearsal of the sleep laboratory increases LD induction beyond a baseline laboratory-related LD induction procedure consisting of explicit sleep lab-related verbal instructions and presleep metacognitive training. We compared this baseline condition with VR rehearsal alone, VR combined with haptic stimulation during REM sleep, and VR containing implicit system anomalies. Lucidity was assessed using both externally rated dream reports and signal-verified eye movements. We hypothesized that additional sensory or contextual manipulations might increase lucidity beyond the baseline context-based induction procedure.

## 2. Methods

### 2.1 General Methods

#### 2.1.1 Participants

All participants reported regular sleep patterns, no history of neurological or psychiatric disorders, and no use of medication known to influence sleep. Informed written consent was obtained from all participants. The protocol was approved by the local ethics committee and conducted in accordance with the Declaration of Helsinki.

#### 2.1.2 Study Design and Procedure

Each session followed the same procedure:

1. Pre-nap preparation: Upon arrival, participants were welcomed and briefed on the session. EEG setup was applied in some cases (see Sleep Recording).
2. VR induction: Participants completed a 10-minute VR task that simulated the lab environment and required them to perform a repetitive task with embedded reality-check prompts. This was followed by a questionnaire targeting the VR experience.
3. Morning Nap: After a guided audio mindfulness session, participants were instructed to go to sleep for approximately a 90-minute nap.
4. Post-nap report: Upon awakening, participants completed a structured dream report and rating scales.

#### 2.1.3 VR Induction Protocol

The VRE was built in Unreal (Epic Games, 2025) and viewed through a Meta Quest 3 headset. The simulation recreated the physical sleep lab setting (Figure 1a) and guided participants through a monotonous task that required them to collect letter objects by exploring the VRE (Figure 1b). After finding a letter, participants were instructed to perform a reality check (counting fingers), reinforcing metacognitive monitoring. The letters could be combined to form a sentence, prompting additional bonus action possible in the VRE: drawing on the walls of the VR sleep lab. The VR scenario was designed to be visually and interactively engaging because of the exploration task. No explicitly surreal or bizarre elements were included; instead, the induction emphasized layered experiences of lab-related reality: the real lab, the virtual lab, and potentially, the dream lab.

**Figure 1:**
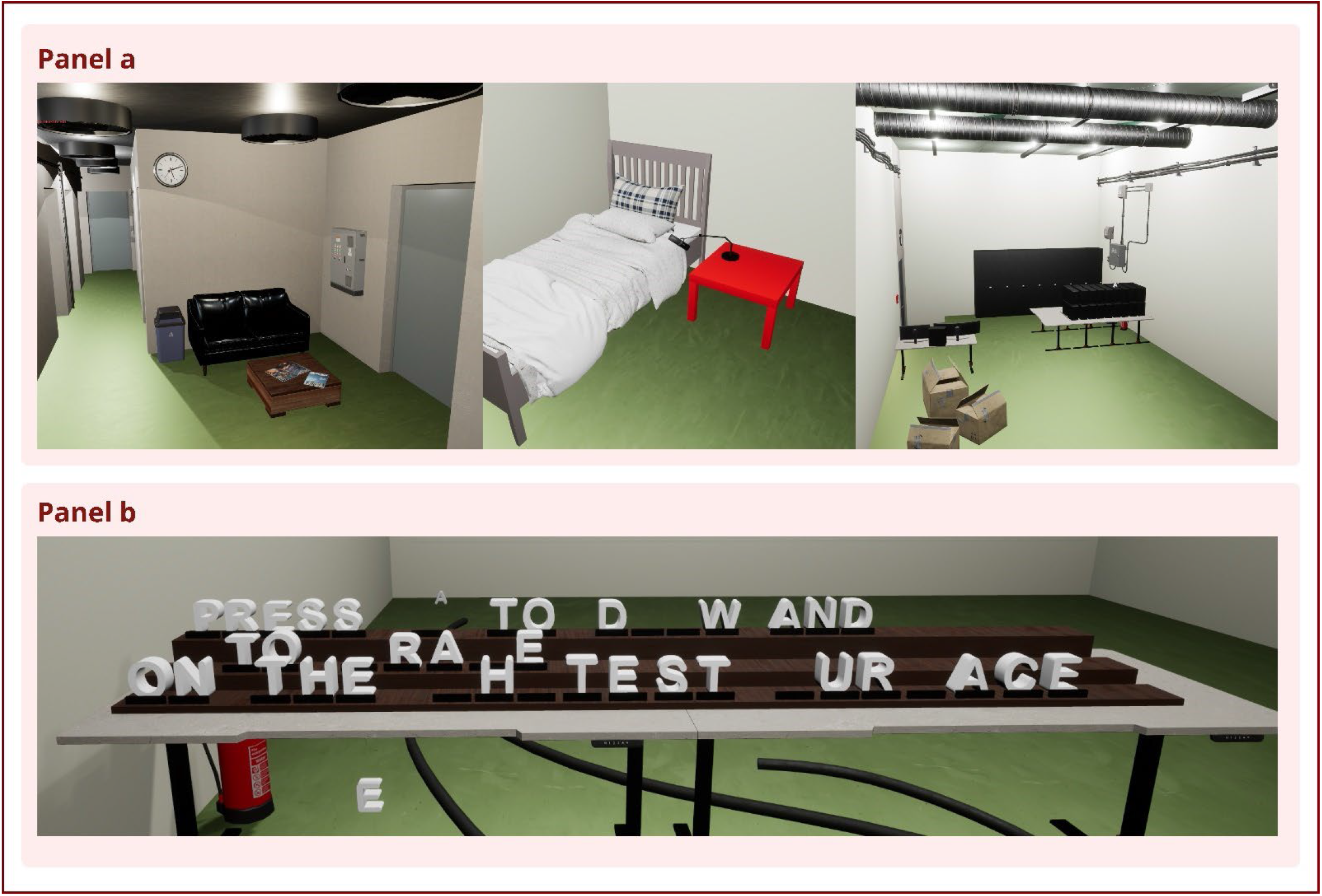
Panel a: Virtual recreation of the sleep lab (in order of appearance: entrance, sleeping room, sensorimotor laboratory. Panel b: VRE-Task; letters could be found within the VRE and arranged to form a sentence, enabling bonus action

#### 2.1.4 Sleep Recording and Dream Reporting

Polysomnography (PSG), including electroencephalography (EEG), electrooculography (EOG), and electromyography (EMG), was utilized to monitor sleep stages during the test naps. PSG setup adhered to the 10-20 system, utilizing 32 active gLADYBIRD electrodes with the gUSBamp EEG system (Klem et al., 1999), using two horizontal EOG, chin EMG, and right earlobe reference. The sampling rate was 256 Hz. Experiment management and streaming were done using the University of Bern’s custom experiment management system Streamix. The sleep room had speakers, a microphone, and a camera for further observation and communication. Before sleep, participants listened to a presleep mindfulness audio guidance based on Carr et al. (2023), focusing on being in the lab and critically reflecting on the situation (see Appendix A). Upon awakening, participants completed an open-ended dream report, followed by the following targeted questions:

1. *‘What went through your head before you woke up?’*
2. *‘Did you notice the stimulation?’*
3. *‘Were you aware that you were dreaming while you were dreaming?’*
4. *‘Did you perform any LRLR eye movements during the dream? ‘*
5. *‘Did you notice anything else?’*

### 2.2 Study Specific Protocols

#### 2.2.1 Study 1 (HS)

Study 1 used a within-subject, counterbalanced design in which 44 participants (N female = 18; M age = 24.1; SD =2.7) completed two morning naps: one following VR training only and one following VR training with additional haptic stimulation (HS). In both sessions, participants completed a 10-minute VR simulation of the sleep lab before the nap. In the HS condition, VR controllers provided vibration during the VR task, and a small tactile device applied brief vibration pulses (2 s) to the left hand during verified REM sleep.

HS was delivered using a custom-built vibrotactile device. The device consisted of a small plastic casing (approximately 6 cm long and 1.5 cm in diameter) containing an eccentric rotating mass (ERM) vibration motor operating at 3 V, with a frequency of 100–150 Hz. The motor was connected to a current-producing LabJack U3‐LV, which allowed the experimenter to trigger the vibration remotely from the control room, producing a light vibration of the plastic tube. The vibrating device was attached to the left dorsal wrist, approximately 2–4 cm proximal to the wrist crease, with its long axis aligned parallel to the forearm and secured using medical adhesive tape. The stimulation was delivered at a single fixed intensity. Prior to lights out, participants briefly experienced the vibration while awake to familiarize themselves with the sensation and to confirm that it was clearly perceptible but comfortable.

REM periods were identified online using the PSG recordings, following AASM criteria. REM onset was marked when low-amplitude mixed-frequency EEG activity, rapid eye movements in the EOG channels, and reduced chin EMG tone were present for at least one 30-second epoch. Only clearly identifiable and stable REM periods were used for stimulation. After REM onset, a timer of 5 minutes was started. Following the 5-minute wait period, a series of 2-second stimulations at a 30-second interstimulus interval (ISI) was presented continuously during REM sleep. If a participant showed signs of arousal or at least 3 epochs of non-REM sleep, stimulation was stopped, and the next REM phase was awaited. No REM awakenings were performed; each nap yielded a single dream report collected at the final awakening. All other procedures, including PSG setup, presleep audio, and dream reporting, followed the general methods.

#### 2.2.2 Study 2 (VR Implicit)

The design of Study 2 was guided by dream reports from the earlier studies, which showed that participants rarely incorporated overtly bizarre elements into their lab dreams. Instead, they consistently described realistic inconsistencies, malfunctioning equipment, unclear procedures, or unexpected experimenter behavior. These subtle, plausible anomalies appeared more likely to influence dream content and occasionally coincided with reflective moments. Based on these observations, we introduced realistic system incongruencies into the VR protocol to mirror the kinds of disruptions that naturally occur in lab dreams and to test whether rehearsing such context-consistent irregularities would facilitate lab incorporation and lucidity more effectively than overt surreal cues.

Study 2 included 38 participants (N female = 27; M age = 31.32, SD = 6.06) recruited online and around the University of Bern campus. Participants completed the same 10-minute VR lab simulation as in Study 1, with intermittent simulated system malfunctions added. During the 10-minute VR session, participants were intermittently exposed to simulated system malfunctions embedded within the virtual environment. These events were presented as full-screen black overlays, displaying short sequences of error messages that lasted approximately 15 seconds each. The messages mimicked realistic device errors to induce brief moments of uncertainty without disrupting immersion. Typical examples included statements such as *“Something went wrong. We’re having trouble completing your request” or “Device disconnected,”* followed by a visible countdown timer that decreased from 10 to 0 seconds. Each sequence concluded with messages such as *“Reconnecting …”* and *“Successfully reconnected.”* The error messages were programmed to appear first 120 seconds after session onset and subsequently at variable intervals between 60 and 100 seconds, looping through the different message types until the session ended. Each interruption temporarily paused the ongoing task and then automatically resumed once the reconnection message was displayed. All other procedures, including nap preparation, polysomnography, and dream reporting, followed the general protocol. No REM awakenings were performed; participants provided a single dream report at the end of the nap to maximize undisturbed REM opportunities.

#### 2.2.1 Study 3 (Active Control)

The active control condition assessed whether explicit lab-focused verbal instructions and presleep audio elicit lucidity. 19 participants (N female = 9; M age = 19.95, SD = 1.54) completed a standard morning nap session with a full polysomnographic (PSG) setup and the same brief presleep audio used in the other studies, which emphasized being present in the sleep lab and reflective awareness. Participants were informed about LD and lab dreaming but received no VR training or sensory stimulation. Unlike Studies 1 and 3, participants were repeatedly awakened during verified REM sleep to obtain immediate dream reports. Each REM period was monitored until signs of arousal or transition to NREM appeared, at which point the participant was awakened and the standard dream report protocol was administered. All other procedures matched the general methods.

### 2.3 Dream Report Analysis

Audio dream reports were transcribed using NoScribe (Dröge, 2025) and subsequently reviewed by a researcher. German dream reports were translated into English by a researcher who spoke both German and English. Dream transcriptions were converted using the dream conversion manual by Schredl (2010). All dream reports were independently evaluated by two raters for lucidity (primary analysis), lab incorporation, FAs, and bizarreness (exploratory analysis) using an adapted version of the *Dream Content Analysis (DCA) Manual* (Schredl, 2010). *Lab incorporation* was coded when the dream narrative contained explicit references to the laboratory environment, equipment (e.g., EEG setup, sensors, computers), or staff members. In cases where only partial elements (e.g., equipment or personnel) appeared without a clear environmental context, raters discussed these instances before reaching a final decision on whether they qualified as full lab incorporation. *Lucidity* was rated based on both explicit statements of dream awareness (e.g., ‘I realized I was dreaming’) and implicit indicators such as volitional control over dream content, deliberate testing of reality, or reflective self-awareness. Additionally, any mention of deliberate left–right–left–right (LRLR) eye movements within the dream was recorded as supplementary evidence of lucidity. These parameters resulted in a single 1 or 0 for lucidity. *False awakenings* were scored as positive when the dream report contained content in which the dreamer woke up in the dream.

### 2.4 Polysomnographic Lucidity Verification

Polysomnographic recordings were exported in HDF5 format and visualized using a custom HDF5 visualizer written in Python. Left-right-left-right eye signals were verified by two individual raters, after which consensus was reached. Figure 2 shows an example of two LRLR eye signals.

**Figure 2:**
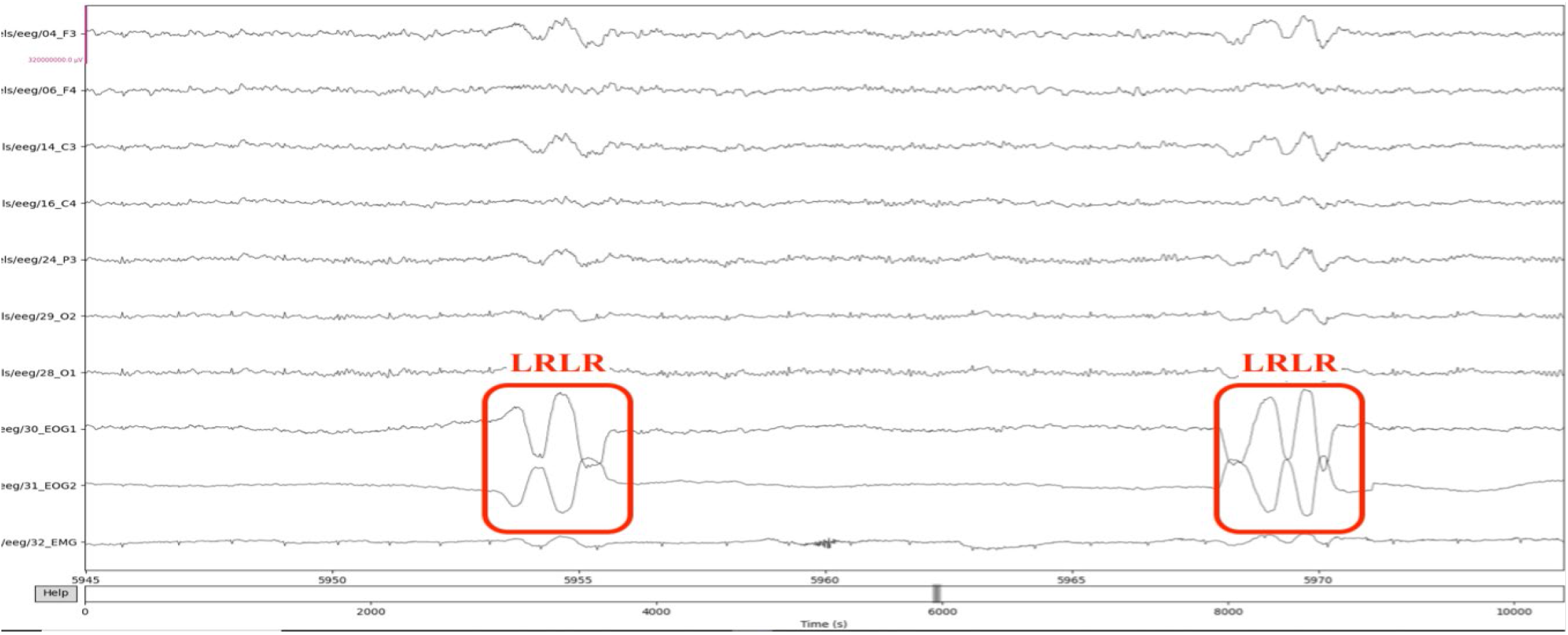
Exemplary LRLR eye movements using electrooculography (EOG) during REM sleep.

### 2.5 Data Selection

The Active Control condition contained multiple dream reports per participant per nap. For the primary analyses, which required one lucid-dream outcome per participant per nap, the dream reports were condensed into one single dream report for each participant. The VR conditions yielded one dream report per nap. For exploratory analyses examining the relationship between lab incorporation and lucidity, all individual dream reports from all studies were included without collapsing across awakenings.

### 2.6 Data Analysis

All statistical analyses were conducted in *Jamovi* (Version 2.6.44, The jamovi project, 2025). The primary dependent variables were LD (0 = no lucid dream; 1 = lucid dream) and Signal-verified lucidity (SVLD), based on LRLR eye movements in the PSG recording (0 = absent; 1 = present). To test whether VR training increased LD relative to the Active Control condition, and whether the VR plus haptic and VR implicit conditions differed from VR alone, we used binomial logistic regression with experimental condition as a fixed factor. These analyses were performed using the GAMLj Generalized Linear Models module with an alpha level of .05.

## 3. Results

### 3.1 Baseline Lucidity

Baseline LD frequency ranged from naïve (never) to frequent lucid dreamers (more than once a month) as displayed in Table 1.

**Table 1:** Lucid dream frequency for each study sample.

| Study | Study 1 | Study 2 | Study 2 | Total |
| --- | --- | --- | --- | --- |
| Lucid Dream Frequency | VR Only and VR + HS (n=44) | Implicit (n=38) | Active control (n=19) | Total (N=101) |
| Frequent: More than 1/Month | 13 (29.5%) | 0 (0%) | 0 (0%) | 13 (12.9%) |
| Rarely: less than 1/month | 16 (36.4%) | 33 (86.8%) | 10 (52.6%) | 59 (58.4%) |
| Never | 15 (34.1%) | 5 (13.2%) | 9 (47.4%) | 29 (28.7%) |

### 3.2 Awakenings and Dream Reports

Across all studies, 161 awakenings were conducted, resulting in 121 dream reports (see Table 2). In the VR conditions (studies 1 and 2), participants were awakened once after the nap, whereas in the Active Control (study 3) participants experienced repeated REM awakenings, yielding multiple reports per participant. For the primary analyses, dream reports in the Active Control group were condensed to a single report per participant, yielding a final sample of 108 dream reports.

**Table 2:** Awakenings and dream reports across studies and conditions.

| Study | Condition | N | Awakenings | Dream Reports (Total) | Dream Reports (Condensed) |
| --- | --- | --- | --- | --- | --- |
| Study 1 | VR Only | 44 | 42* | 29 | 28 |
|  | VR+HS | 44 | 42* | 27 | 25 |
|  | **Total** | 44 | 84 | 56 | 53 |
| Study 2 | Implicit | 38 | 38 | 37 | 37 |
| Study 3 | Active Control | 19 | 39** | 31 | 18 |
| **Total** |  | 101 | 161 | 121 | 108 |

Note: \*Four participants from study 1 only completed one of the conditions, resulting in 42 awakenings for each condition. \*\* Study 3 conducted REM awakenings, resulting in multiple awakenings and dream reports per participant.

### 3.1 Lucidity

Lucidity was analyzed using the condensed dataset (N = 108 dreams). Overall, 45.4% of dreams were externally rated as lucid. A GLMM revealed no significant effect of condition, χ^2^(3) = 0.91, p = .823 (see Table 3). Signal-verified lucid dreams (SVLDs), confirmed via LRLR eye movements in PSG, showed no significant differences between conditions, χ^2^(3) = 3.75, p = .289 (see Table 3). One VR+HS case could not be verified due to technical issues and was excluded.

**Table 3:** Lucidity outcomes across studies and conditions with GLMM results.

| Outcome | Condition | n Lucid / N (%) | Omnibus GLMM $\chi^2$(df), p |
| --- | --- | --- | --- |
| **Externally Rated LD** | VR Only | 12 / 28 (42.9%) | $\chi^2(3)=0.91$, p=.823 |
|  | VR + HS | 11 / 25 (44.0%) |  |
|  | Active Control | 7 / 18 (39.9%) |  |
|  | Implicit | 19 / 37 (51.4%) |  |
| **Signal-Verified LD (SVLD)** | VR Only | 9 / 28 (32.1%) | $\chi^2(3)=3.75$, p=.289 |
|  | VR + HS | 4 / 25 (16.0%) |  |
|  | Active Control | 2 / 18 (11.1%) |  |
|  | Implicit | 11 / 37 (29.7%) |  |

#### Exploratory analysis: Lab dreaming, false awakenings, and bizarreness

All 121 dream reports were included in this exploratory analysis. Laboratory-related content was present in 58 dreams (47.9%), false awakenings (FA) occurred in 28 dreams (23.1%), and lucid awareness was reported in 52 dreams (43.0%). As illustrated in Figure 3, substantial overlap was observed between these categories. Notably, 17 dreams involved lucidity emerging after a false awakening within the laboratory context. Within laboratory dreams, most reports referenced the laboratory environment (91.4%), while lab-related characters (60.3%) and laboratory equipment (36.2%) were mentioned less consistently.

**Figure 3:**
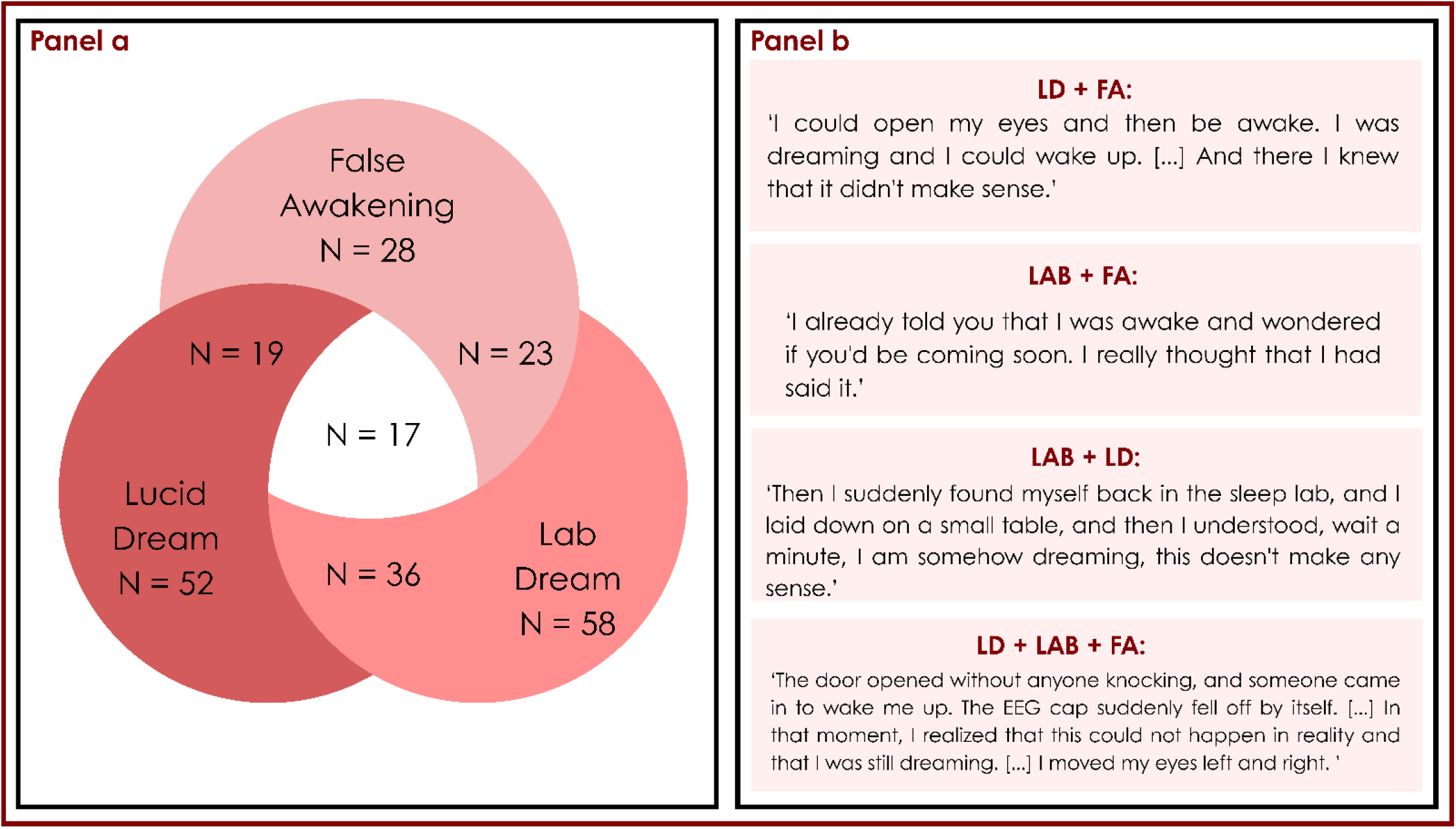
Panel a: lucid dreaming, false awakenings, and lab dreaming co-occurrence. Panel b: excerpts illustrating the co-occurrence of dream categories.

## Discussion

The present series of studies tested whether LD induction can be enhanced by strengthening waking-dream continuity through immersive presleep VR rehearsal in the sleep laboratory and whether additional haptic stimulation or implicit VR anomalies further modulate lucidity. Participants in all conditions received explicit lab-related LD-induction instructions and presleep training to target recognition of the laboratory environment as a dream sign. A secondary exploratory analysis examined the overlap between laboratory dreaming, FAs, and lucidity. The present work is therefore a systematic test of context-based LD induction in which participants were explicitly trained to recognize a predictable dream environment.

Across conditions, externally rated lucidity occurred in approximately 43–51% of dreams, and signal-verified lucid dreams occurred in 11–32% of REM periods. These rates are comparatively high for a single laboratory nap in a largely non-expert sample, suggesting that the combination of explicit lucidity instructions and the salience of the laboratory environment may have created favorable conditions for lucidity. Using this design, immersive VR rehearsal of the laboratory environment did not increase LD induction beyond the baseline laboratory-based induction procedure. Similarly, neither the addition of haptic stimulation nor the introduction of subtle VR anomalies significantly altered the likelihood of lucidity. Although statistical comparisons revealed no significant differences between conditions, some numerical variation was observed (Table 3). For example, signal-verified lucid dreams occurred in 32.1% of participants in the VR-only condition and 29.7% in the implicit VR condition, compared with 16.0% in the VR + HS condition and 11.1% in the Active Control condition. Given the relatively small number of signal-verified events, these differences should be interpreted cautiously and may reflect sampling variability rather than systematic condition effects. In previous control conditions without any LD-related instructions, success rates of 9% and 0% were observed (Erlacher & Stumbrys, 2020). Our findings suggest that factors shared across conditions, particularly explicit lucidity instructions and the salience of the sleep-laboratory environment, may have played a stronger role in promoting lucidity than the specific experimental manipulations. The sleep-laboratory context itself reliably elicits congruent dream content (Picard-Deland et al., 2021; Schredl, 2008), and explicit training focused on recognizing this environment may therefore provide a natural pathway to lucid awareness. VR training did not outperform immersion in a LD-focused laboratory setting. Haptic stimulation, designed to cue reality checking during REM sleep physically, did not further enhance lucidity, suggesting that bottom-up tactile cues may be insufficient to alter metacognitive dynamics during REM sleep (Peters et al., 2024). Similarly, subtle system incongruities within the VRE did not yield higher lucidity rates than in the other conditions.

The exploratory analyses revealed substantial overlap between laboratory dreams, FAs, and LDs. Lucid awareness frequently emerged following a FA within the laboratory context, consistent with longstanding observations linking FAs to lucidity (Buzzi, 2011, 2019; Hearne, 1982) (Figure 3). FAs, highly realistic dreams of awakening, can precede or terminate lucidity and may occur in loops (LaBerge & DeGracia, 2000). Reality checking has repeatedly been described as the mechanism by which FAs transition into lucid dreams (Buzzi, 2019; Hearne, 1982), precisely the strategy practiced in the present study. The data, therefore, strengthen the view that false awakenings serve as a natural gateway to metacognitive activation within realistic dream environments.

Within predictive-processing accounts of dreaming, lucidity has been thought to emerge when prediction errors cannot be seamlessly integrated into the ongoing narrative (Hobson et al., 2014; Hobson & Friston, 2012). Our data do not support causal claims regarding the specific contribution of VR to reality checking or prediction-error detection. Instead, the findings highlight the laboratory environment itself as a cue that may lead to reflective awareness. The implicit VR manipulation introduced minor system-error messages that resembled the procedural errors often reported in laboratory dreams. While this condition did not increase lucidity, its conceptual alignment with models of moderate prediction conflict remains theoretically relevant. Given that emotionally salient experiences are more likely to be incorporated into dreams (Schredl, 2008), even mild uncertainty during VR exposure could, in principle, influence subsequent dream content. However, these theoretical interpretations remain speculative, as the present study did not assess metacognitive monitoring, emotional salience during VR, or prediction errors directly. Previous VR-based research has reported increased lucidity following repeated exposure to surreal VR environments (Gott et al., 2020). However, effects were observed relative to the passive but not the active control conditions during practice of standard induction techniques. The present findings are consistent with this pattern: lucid dreams occurred frequently across all conditions, suggesting that explicit induction training within the laboratory context may already be a strong driver of lucidity.

Several limitations should be considered. The control condition differed procedurally from the VR conditions in that repeated REM awakenings were performed, potentially affecting opportunities for recall and the probability of lucidity. Although dream reports were condensed for primary analyses, this structural difference limits strict causal comparison between conditions. In addition, the brief 10-minute presleep training may have been insufficient to detect learning-dependent effects, as many established induction techniques rely on repeated exposure across multiple nights. Participants were also explicitly informed about lucid dreaming and received lucidity-focused instructions, which may have elevated baseline lucidity across all conditions. In addition, the relatively small number of signal-verified lucid dreams limits statistical power to detect small differences between conditions.

Another limitation concerns the ecological validity of the paradigm. The context-based induction strategy relied on the salience of the sleep-laboratory environment, which participants had experienced immediately prior to sleep. Because laboratory settings are unusual and highly structured, they may be especially likely to appear in dreams and prompt reflective awareness. As a result, the current approach may be less directly applicable outside laboratory studies where such a shared context is absent.

Overall, the results indicate that LD induction in this study was influenced more by contextual and cognitive factors than by specific technological manipulations. Lucidity often emerged in realistic laboratory dream scenarios, particularly after FAs, suggesting that moments of ambiguity in familiar dream environments may naturally trigger reflective awareness. Although VR provided a controlled way to rehearse the sleep-laboratory context, it did not increase lucidity beyond the baseline context-based induction procedure.

More broadly, these findings highlight the potential of context-based approaches to LD induction. Rather than relying solely on sensory cues during sleep, induction strategies may benefit from targeting dream environments likely to recur. Future research could extend this approach beyond the laboratory by using immersive simulations to train recognition of common dream scenarios. For example, VREs could recreate frequently reported dream settings such as returning to school, being at work, or navigating unfamiliar buildings. Repeated exposure to such scenarios may allow individuals to practice linking these contexts to reality-checking or reflective awareness. If these trained situations subsequently appear in dreams, they may serve as triggers for lucidity. Testing such paradigms across repeated training sessions and in home-sleep settings would help determine whether VR can support scalable context-based lucid-dream induction outside the laboratory. These findings suggest that leveraging predictable dream contexts may represent a promising direction for LD induction research, complementing traditional cue-based approaches that attempt to influence dreams from within sleep itself.

## Acknowledgements

We would like to extend our gratitude to Tobias Aubert for the VR programming, Lukas Schmid and Susanne Zulauf for data acquisition, and the Technology Platform for Science (TPF) of the Faculty of Human Sciences of the University of Bern.

## Appendices

### Appendix A: Presleep Targeted Lucidity Reactivation Audio Training. Total length: 22:34min

We are now going to practice becoming lucid. We want to train your mind to recognize the lab as a lucidity cue, so that you can have a lucid dream. Every time you think about the sleep lab, reflect on what you’re experiencing in that moment. You will become lucid by attending to where your mind has been, attending to your body, and attending to your surroundings. Focus on how aspects of your experience might be in any way different from your normal waking experience. You do not need to do the eye signals while awake. Only do the signals when you become lucid in a dream. Start by closing your eyes and taking a few deep breaths, inhaling deeply through the nose and exhaling slowly through the mouth. Allow each breath to relax you more deeply, letting go of any tension or stress you may be holding on to.

As you lie here in this bed, you become lucid. Bring your attention to your thoughts and notice how your mind has wandered. Now observe your body, sensations, and feelings. Observe your breathing, and remain critically aware, lucid, and notice how aspects of this experience are in any way different from your normal waking experience.

Repeat four times

Remember, each time you think about the lab, practice becoming lucid by observing your thoughts, body, and feelings. And notice how aspects of this experience differ in any way from your normal waking experience.

You can now begin to fall asleep. Keep trying to become lucid by focusing on how your mind has wandered. Observe your thoughts, body, and feelings. Remember to perform a left-right, left-right, center eye-movement when you become lucid in a dream.

